# Refactored genetic parts for modular assembly of the *E. coli* MccV type I secretion system used to screen class II microcin candidates from plant-associated bacteria

**DOI:** 10.64898/2026.01.19.700402

**Authors:** Alexa K. Morton, Krisha Chaudhari, Brian D. Matibag, Vibhav B. Iyengar, Katherine E. Dullen, Anthony J. VanDieren, Jennifer K. Parker, Dennis M. Mishler, Jeffrey E. Barrick

## Abstract

**Background:** Microcins are small antibacterial proteins secreted by gram-negative bacteria. The activities of new microcins discovered using bioinformatic searches need to be validated and characterized to understand how they mediate competition in microbiomes and to evaluate their potential as new therapeutics for combating antibiotic resistance. Engineered plasmids containing the type I secretion system associated with *Escherichia coli* Microcin V (MccV) can secrete heterologous proteins, including other class II microcins, and this system functions in other bacterial hosts. However, existing microcin secretion constructs are not designed for easily swapping components — such as origins of replication, resistance genes, promoters, and signal peptides — that may need to be changed for compatibility with other chassis.

**Results:** We refactored the *E. coli* MccV type I secretion system into genetic parts compatible with a modular Golden Gate assembly scheme and used these parts to construct two-plasmid microcin secretion systems. In our design, one plasmid encodes the type I secretion system proteins, and the other encodes a signal peptide fused to the cargo protein that will be secreted. We tested two versions of a system with inducible promoters separately controlling expression of the components on each plasmid. One used plasmids that replicate in *E. coli* and its close relatives. The other used broad-host-range plasmids. When induced to secrete MccV, both versions produced similar zones of inhibition against a susceptible strain of *E. coli*. Next, we identified putative class II microcins in genomes of bacteria from plant-associated genera (*Pantoea*, *Erwinia*, and *Xanthomonas*) using an existing bioinformatics pipeline. We screened 23 of these putative microcins for *E. coli* self-inhibition. Seven exhibited some inhibition, mostly later in growth curves, but none had effects that were comparable in strength to MccV.

**Conclusions:** The genetic parts we created can be assembled in various combinations into tailored systems for secreting small proteins from diverse bacterial chassis. These systems can be used to further characterize the targets of novel microcins and to secrete these or other small proteins for various applications. For example, beneficial bacteria used in crop protection could be engineered to secrete microcins that kill or inhibit plant pathogens to increase their efficacy.

## Background

Microcins are an emerging class of antibacterial proteins secreted by gram-negative bacteria (1–5). In nature, microcins mediate microbial competition, often by targeting bacteria that are closely related to the microcin-producing strain (6–8). This narrow activity spectrum makes microcins an appealing alternative to broad-spectrum antibiotics in terms of managing the evolution of resistance. Class II microcins are translated by the ribosome and undergo no posttranslational modifications except the formation of disulfide bonds (class IIa) or the addition of a C-terminal siderophore (class IIb) (3). The relative simplicity of class IIa microcins makes these potential therapeutics especially amenable to expression in heterologous hosts (2, 9).

Class II microcins are secreted from bacterial cells by a specialized type I secretion system (T1SS) consisting of a C39 peptidase domain-containing ABC transporter (PCAT) and a membrane fusion protein (MFP) that interact with the TolC outer membrane channel protein that is shared with other efflux systems (1, 9, 10). Class II microcins are translated with an N-terminal signal peptide, which is recognized by the PCAT and cleaved during export to produce the mature microcin. Microcin-producing bacteria also typically express a cognate immunity protein that prevents self-killing. PCAT, MFP, microcin, and immunity proteins are often encoded together in a genome, which has aided bioinformatic searches for new class II microcins (2, 11).

The T1SS of the *E. coli* class IIa Microcin V (MccV), has been cloned into one- and two-plasmid systems, engineered to secrete heterologous microcins and small proteins (9, 12, 13), and shown to function in other bacterial hosts (2), including *Vibrio cholerae* (9). We wanted to streamline the process by which one can use this T1SS to test newly identified microcins for activity and secrete other small proteins for various applications. To do so, we converted the MccV T1SS components into genetic parts compatible with an existing Golden Gate assembly scheme (14, 15). This refactoring makes it possible to rapidly construct plasmids with different combinations of promoters, origins of replication, resistance genes, secreted protein cargos, and microcin T1SSs from species other than *E. coli*. We validated that two-plasmid systems constructed from these parts with inducible promoters controlling T1SS and microcin expression were functional.

MccV and other class II microcins from *E. coli* and related bacteria in the *Enterobacteriaceae* are the best-studied examples of this class (6). However, class II microcins have been discovered beyond this family (2, 4, 5, 16), and recently many more microcins with associated T1SSs have been predicted using new bioinformatic tools (2, 11, 17). These developments inspired us to search for class II microcins in the genomes of *Erwinia, Pantoea*, and *Xanthomonas* species. These genera include important plant pathogens and also strains that are used as biocontrol agents to protect plants from pathogens (18–21). We identified many putative class II microcins alongside T1SS components within these bacterial groups. As a first step in characterizing their functions, we used our modular MccV T1SS plasmids to screen 23 of these putative class IIa microcins for activity against *E. coli*.

## Methods

### Culture conditions

*E. coli* were incubated at 37 °C for growth. Liquid cultures were grown in test tubes with 200 r.p.m. orbital shaking over a 1-inch diameter. The Miller formulation of Lysogeny Broth (LB) (10 g tryptone, 5 g yeast extract, 10 g NaCl per liter) and M9 minimal medium (3 g KH_2_PO_4_, 6.8 g Na_2_HPO_4_, 0.5 g NaCl, 1 g NH_4_Cl per liter) supplemented with 0.4% (v/v) glycerol (M9-glycerol) were used as growth media. SOC medium was used for transformation (20 g tryptone, 5 g yeast extract, 0.5 g NaCl, 0.186 g KCl, 2.4 g MgSO_4_ per liter) supplemented with 0.4% (v/v) glucose. LB plates included 16 g/L agar and 0.005% (v/v) antifoam B emulsion (Sigma-Aldrich). When applicable, media were supplemented with antibiotics at the following concentrations: 20 µg/mL chloramphenicol (cam), 50 µg/mL kanamycin (kan), and 100 µg/mL carbenicillin (carb). Inducers were added to media at final concentrations of 10 µM for 3-hydroxytetradecanoyl-homoserine lactone (OHC14) and 100 µM for vanillate. Strains used in this work are described in **Table S1**. Bacterial cell stocks were stored at −80 °C in 15–20% (v/v) glycerol.

### Plasmid assembly

We used the Golden Gate assembly (GGA) scheme from the Bee Microbiome Toolkit (BTK) and its progenitor, the Yeast Toolkit (YTK), to construct our plasmids (14, 15). NEBridge BsaI and BsmBI Golden Gate Assembly kits (New England Biolabs) were used for all GGA reactions. Part plasmids were assembled using the *camR* GFP dropout entry vector pBTK1001 to insert either PCR products or synthetic dsDNA fragments via BsmBI assembly, and transformant colonies were selected by screening for GFP-negative colonies on LB+cam. Part plasmids were further assembled into mScarlet dropout vectors (pBTK1028, pBTK1063, pBTK1065, or pBTK1079) using GGA with BsaI restriction sites to create Stage 1 assembly plasmids. These assembly plasmids contain either a *kanR* or *ampR* gene, and transformant colonies were selected by screening for mScarlet-negative colonies on LB+carb or LB+kan. Transformations were conducted via heat shock at 42 °C into chemically competent *E. coli* DH5α. After transformation, cells were incubated in 1 mL of SOC for 1 h prior to selective plating. Plasmids were isolated from transformed cells using QIAPrep Spin Miniprep Kits (QIAGEN). All part plasmids were verified using Oxford nanopore sequencing (Plasmidsaurus). Plasmids originating in this work are described in **Table S2**, and their full DNA sequences are provided in **Data S1**.

### T1SS refactoring and domestication

Our designs built on previously published MccV T1SS plasmids (9). This previous secretion system has two versions: (1) a single plasmid system (pMMB67EH encoding PCAT, MFP, and cargo) and (2) a two-plasmid system (pACYC184 encoding PCAT and MFP + pBAD18 encoding cargo). We rearranged and modified genetic parts from these plasmids to make them compatible with GGA following the BTK/YTK syntax (14, 15). **Table S3** contains the relevant BsaI assembly overhang sequences used to join parts. We used plasmid pSK03 as a template to amplify parts encoding the MccV immunity protein (*cvi*), the bicistronic MFP (*cvaA*) and PCAT (*cvaB*) operon (*cvaAB*), the MccV signal peptide (SP), and the mature MccV microcin (*mccV*). The SP and *mccV* together comprise the MccV premicrocin (*cvaC*). As in the original plasmid systems, we did not clone TolC since it is a general secretion system factor that is widespread in gram-negative bacteria, even those that do not natively encode microcin T1SSs (10).

Our assembly scheme divides the reading frame (a Type 3 BTK part) for the secreted protein cargo into Type 3p and Type 3q parts encoding the signal peptide and cargo respectively. In the final construct, these two parts are fused in-frame through a new TGGC overhang that overlaps with the double-glycine motif at the SP cleavage site. When expressing a microcin, the immunity protein can be included downstream as part of the same transcriptional unit. In the case of *cvi* and *cvaC*, this change required switching the order of these genes from the natural configuration (in which *cvaC* is located downstream of *cvi*) and removing 23 bp of overlap between their reading frames. We domesticated the Type 3 *cvaAB* part sequence by making a synonymous point mutation in *cvaB* to remove an internal BsaI restriction site.

For implementing inducible expression, we refactored systems from the Marionette YFP sensor plasmids (22) into Type 2 parts containing inducible promoter sequences and ribosome binding sites and Type 4 parts that express the associated transcriptional regulators. These Type 4 parts include an upstream terminator to insulate expression from the upstream transcriptional unit.

### Inducible promoter function

We assembled plasmids with our inducible promoter system parts controlling expression of *gfpmut3* (14). To test induction, we revived *E. coli* DH5α strains with these plasmids from glycerol stocks by inoculating 5 mL of LB containing appropriate antibiotics. After overnight growth, we transferred 5 µL from these cultures into another 5 mL of LB with antibiotics. At this stage, we inoculated five replicates with and five replicates without the small molecule inducer for each construct. These cultures were grown overnight alongside an uninoculated LB blank and a culture of DH5α containing no plasmid in LB without antibiotics. We transferred 300 µL from each induced and uninduced culture into a black clear-bottom 96-well microplate alongside five wells filled from the LB blank and five wells filled with the no plasmid control cultures. We loaded this microplate into a Tecan Infinite Pro M200 Plate Reader and measured GFP fluorescence (λ_ex_ = 485 nm, λ_em_ = 535 nm) and optical density at 600 nm (OD600) for each well.

To process the resulting data, we first calculated the mean OD600 and mean relative fluorescence unit (RFU) values of the LB blank wells. We subtracted these background values from the OD600 and RFU values of all other wells. The RFU value of each well was then normalized by dividing by its OD600. Next, we calculated the mean of the normalized RFU values of the no plasmid control wells. Finally, we subtracted this value from the normalized RFU values of all wells containing engineered plasmids to produce the final RFU values for the amount of fluorescence produced per cell under induced and noninduced conditions.

### Zone of inhibition assays

To assess the effectiveness of the refactored MccV secretion system, we performed zone of inhibition (ZOI) assays against *E. coli* W3110 which is susceptible to killing by this microcin. Solid M9-glycerol supplemented with 2 mM MgSO_4_ and 0.1 mM CaCl_2_ was used as the growth medium in these assays. Plates were poured with a thin base layer containing 15 g/L agar. To begin each assay, *E. coli* W3110 and the strains to be tested for MccV secretion were revived from glycerol stocks in 5 mL LB cultures with appropriate antibiotics. After overnight growth, we transferred 5 µL into 5 mL LB cultures that included appropriate inducer molecules and excluded antibiotics to avoid carryover into the assay. After overnight growth again, we took OD600 readings. We concentrated cultures of the strains being tested for microcin secretion by centrifuging them at 4500 × g for 1 min. Then, we removed the supernatant and resuspended these cultures in saline at an OD600 of 50. We inoculated melted media containing 7.5 g/L agar and the appropriate inducers with W3110 cells at an OD600 of 0.01 and poured 5 mL on each plate containing a base layer of agar. After the top agar solidified, 10 µL of each strain being tested for secretion was spotted on top. All plates included spots of *E. coli* DH5α with no plasmids as a negative control and *E. coli* W3110 strain SK01, which is known to secrete MccV effectively (9), as a positive control. Plates were photographed after incubation overnight.

### Microcin candidate identification and cloning

We used *cinful* v1.2.6 (11) to identify putative class II microcins in the genomes of *Pantoea*, *Erwinia*, and *Xanthomonas* strains downloaded from GenBank. Microcin candidates were filtered to those found by a hidden Markov model trained on known microcin sequences (hmmerHit = TRUE) with T1SS components encoded in the same genome (MFP and PCAT proteins identified). Next, we aligned predicted microcin sequences using MUSCLE v5.3 (23) to identify N-terminal signal peptides of 15-20 amino acids ending with an expected GG, GA, or GS cleavage site motif (24, 25). Through manual inspection, we further eliminated microcin candidates that did not have plausible signal peptides and those with C-terminal Gly/Ser repeats. The latter are often associated with siderophore modifications in class IIb microcins (3). The full list of putative class II microcins passing these criteria is provided in **Table S4**.

Of the remaining candidates, we selected the ones to clone and test as follows. We prioritized those with T1SS components or candidate immunity proteins (hypothetical proteins or unannotated reading frames encoding 50–250 amino acid proteins) encoded nearby in the genome. We used FastTree v2.1.11 (26) on a MUSCLE alignment of microcin sequences (with signal peptides omitted) to cluster them so that we could select representatives from different sequence families. Five known class IIa microcins and five known class IIb microcins were included (1). Unrooted trees of all final microcin candidates divided by genus can be found in **Figure S1**. Full *cinful* results from before filtering and passing microcin candidate lists, sequence alignments, and phylogenetic trees are provided in text file formats in **Data S2**.

DNA sequences encoding microcin candidates (omitting predicted signal peptides) were extracted from genome sequences. We ordered these as synthetic double-stranded DNA fragments flanked with BsmBI and BsaI restriction sites and overhangs for cloning them into an entry vector to make Type 3q part plasmids, as described above. *Pantoea* microcin candidates were ordered as gBlocks from Integrated DNA Technologies. *Erwinia* and *Xanthomonas* microcins were ordered as gene fragments from Twist Biosciences.

### *E. coli* self-inhibition assays

We transformed *E. coli* W3110 with the secretion system plasmid. Then, we transformed these cells with each cargo plasmid expressing a microcin candidate, a positive control version of the cargo plasmid expressing MccV, or a negative control version expressing the synthetic peptide G3P2, which has a randomized amino acid sequence that is not toxic to *E. coli* (9).

To begin each assay, we streaked glycerol stocks of each strain being tested on LB+kan+carb agar and incubated these plates overnight. Then, we picked separate colonies to inoculate eight LB+kan+carb liquid cultures of each strain. After overnight growth, cultures were diluted to an OD600 of 1.0 in M9-glycerol supplemented with 0.2% (w/v) casamino acids, kan, and carb. Next, we prepared 200 μL duplicates of each culture, one induced and the other uninduced, in a clear, flat-bottom 96-well plate. To do so, we added 20 μL of each diluted culture to 180 μL of M9-glycerol with the same amendments, and, if induced, also included OHC14. The 96-well plate was then loaded into a Tecan Infinite Pro M200 Plate Reader and incubated at 37 °C with orbital shaking. OD600 values were measured every 15 min for 12 h.

We tested each of the 23 microcin candidates and the two controls in two separate experiments. The resulting set of 50 total growth curves was analyzed for inhibition at the 4 h and 12 h timepoints. To determine whether inhibition was statistically significant, we first required that the mean OD600 of the induced cultures be ≤ 90% that of the mean OD600 of the uninduced cultures at a timepoint. Second, we performed one-tailed *t*-tests on the OD600 measurements with and without induction and corrected the *p*-values using the Benjamini-Hochberg method with a 5% false-discovery rate within each set of 50 pairs of measurements at a time point. We required that the adjusted *p* value be < 0.05 for inhibition to be judged as significant.

## Results

### Design and cloning of modular microcin secretion system parts

We designed a modular cloning scheme that can incorporate different combinations of T1SS components, signal peptides, secreted cargo proteins, immunity proteins, and inducible promoter systems into DNA constructs (**Fig. 1**). We implemented this scheme by refactoring sequences from an existing *E. coli* MccV T1SS plasmid (pSK03) (9) into genetic parts compatible with the Golden Gate assembly standards used by the BTK and YTK toolkits (14, 15). In these toolkits, Stage 1 assembly of part Types 1-8 through compatible BsaI overhangs creates a complete plasmid. By convention, Type 2 parts contain promoters and ribosome binding sites, Type 3 parts are individual open reading frames or operons, and Type 4 parts are terminators. To simplify assembly, we consolidated Type 5, 6, 7, 8, and 1 parts into a series of Type 56781 dropout vectors with different antibiotic resistance genes paired with either narrow-host-range (NHR) or broad-host-range (BHR) origins of replication. These dropout vectors were used as backbones for Stage 1 assembly to construct all plasmids we tested.

**Fig. 1:**
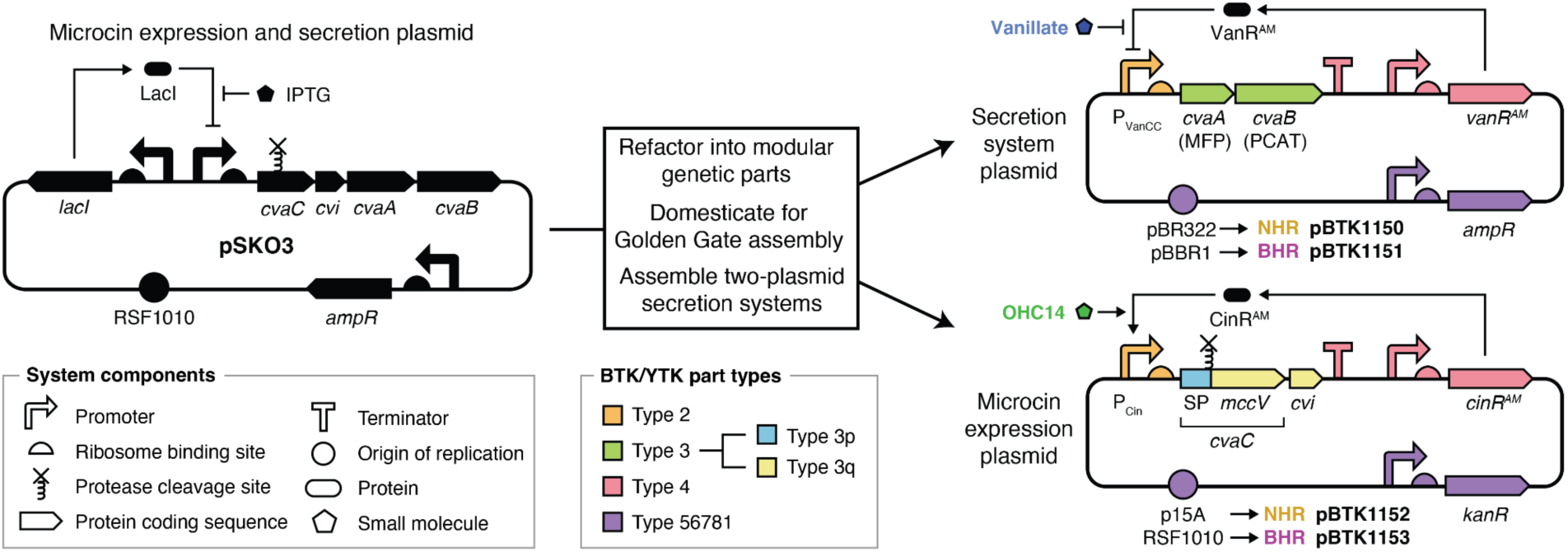
Refactoring the *E. coli* MccV T1SS system into genetic parts for modular assembly. T1SS components and the microcin cargo from plasmid (pSK03) were domesticated and cloned into part plasmids for Golden Gate assembly according to the BTK/YTK scheme. We constructed two-plasmid secretion systems from these parts. The secretion system plasmid encodes the CvaA (MFP) and CvaB (PCAT) components of the T1SS, which function together with TolC natively encoded by the host to secrete proteins tagged at the N-terminus with the cognate signal peptide (SP). The microcin expression plasmid expresses the cargo protein targeted for secretion fused to the SP. In the example shown, the cargo is *E. coli* Microcin V (MccV), which is produced from the pre-microcin CvaC when the PCAT removes the SP during secretion. The Cvi immunity protein that protects the microcin-secreting cell from MccV is expressed as part of the same operon. Orthogonal promoter/regulator pairs P_VanCC_/VanR^AM^ and P_Cin_/CinR^AM^ control expression of the secretion system and the secreted protein cargo (plus immunity gene, in this case). BTK/YTK part modules are color coded by their types defined by the BsaI overhangs used for Stage 1 assembly. We split the Type 3 module in the cargo plasmid into 3p/3q subparts by defining a new overhang between them that overlaps the SP diglycine cleavage site. Different origins of replication were used in the narrow-host range (NHR) and broad-host-range (BHR) versions of the two-plasmid system.

To adapt the *E. coli* MccV T1SS for the BTK/YTK assembly scheme, we first created a Type 3 domesticated MFP and PCAT operon part with a silent mutation that eliminated a BsaI restriction site. Then, we defined a new BsaI overhang that was orthogonal to the existing ones and used it to split a Type 3 part into 3p and 3q subparts. This refactoring makes it possible to fuse together different secretion tag and cargo protein parts into one open reading frame. We created a Type 3p part that encodes the signal peptide from the MccV premicrocin, but the new overhang is compatible with any signal peptide with a diglycine (GG) at the cleavage site. Type 3q parts encode the mature microcin or other small protein cargo targeted for secretion. We created Type 3q parts encoding MccV and MccV plus its cognate immunity protein.

### Refactored inducible promoter parts maintain performance

In order to separately control expression of the T1SS and its secreted protein cargo, we refactored the VanR and CinR inducible promoter systems from the *E. coli* Marionette sensor collection (22) to make them compatible with the BTK/YTK assembly scheme. We selected these inducible promoter systems because they have been shown to function not only in *E. coli* but also in a variety of Proteobacteria (27). For each system, we created a Type 2 part containing the regulated promoter and a ribosome binding site plus a Type 4 part that constitutively expresses the regulatory gene with an upstream terminator for insulation (**Fig. 1**).

To test the performance of the refactored parts, we cloned *gfpmut3* under control of each inducible promoter system into the NHR p15A backbone and transformed these plasmids (**Fig. 2a**). For *E. coli* DH5α transformed with these plasmids, we observed similar performance to what has been reported in prior studies (22, 27), with >50-fold higher GFP signal in induced cultures (**Fig. 2b**). To test performance in the BHR framework, we cloned *gfpmut3* under VanR control on a pBBR1 plasmid and under CinR control on an RSF1010 plasmid and transformed these plasmids (**Fig. 2a**). When these BHR constructs were tested in *E. coli* DH5α, we observed similar performance to the NHR constructs. With this proof of performance in *E. coli*, we incorporated the refactored Marionette systems into our microcin secretion plasmids.

**Fig. 2:**
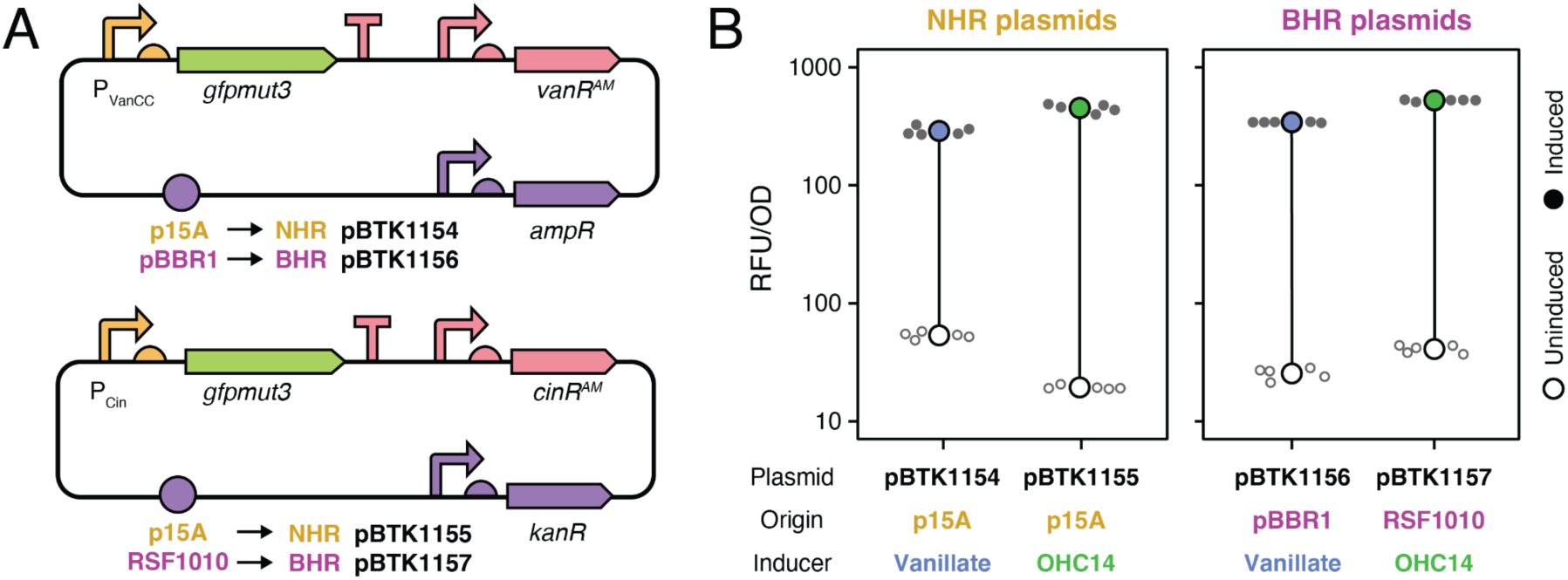
Refactored VanR and CinR inducible promoter systems function in *E. coli*. **(A)** Plasmids constructed to test inducible promoter parts. The *gfpmut3* gene was cloned under control of promoter/regulator pair P_VanCC_/VanR^AM^ (induced by vanillate) or P_Cin_/CinR^AM^ (induced by OHC14). Both systems were tested on narrow-host-range (NHR) and broad-host-range (BHR) plasmid backbones. Constructs are colored by part types according to the legend in Figure 1. **(B)** GFP signal from cells containing inducible promoter plasmids in relative fluorescence units (RFU) normalized by optical density at 600 nm (OD600). Mean values are shown for induced (filled circles) and uninduced (open circles) cultures alongside the individual values for each of five biological replicates (smaller circles).

### MccV secretion system plasmid assemblies function in *E. coli*

To validate that our refactored genetic parts functioned, we assembled the components of the *E. coli* MccV T1SS into two-plasmid systems for testing (**Fig. 1**). One plasmid encodes the machinery necessary to export class II microcins from gram-negative bacteria, the MFP and PCAT. As was the case for its predecessors, this system relies on TolC expression by the host to complete the T1SS. The other plasmid contains the secreted protein cargo fused to the MccV signal peptide. We tested two-plasmid systems engineered to secrete MccV that encode its associated immunity protein as part of the same operon. We constructed a NHR version of this two-plasmid system using the p15A and pBR322 origins of replication that function in *E. coli* and its close relatives (28, 29) and a BHR version using the pBBR1 and RSF1010 origins of replication that replicate in more diverse bacterial hosts (30).

We transformed these NHR and BHR two-plasmid MccV secretion systems into *E. coli* DH5α and tested whether MccV secretion led to a zone of inhibition when these strains were spotted on top agar containing *E. coli* W3110, which is susceptible to MccV killing. Both the NHR and BHR sets of plasmids use VanR to control expression of T1SS components and CinR to control expression of MccV. We observed a strong zone of inhibition when MccV expression was induced with OHC14, whether or not T1SS expression was induced with vanillate (**Fig. 3**). This result suggests that very little MFP and PCAT expression is necessary for effective secretion and that enough of these components are produced by uninduced (leaky) expression from the VanR-regulated promoter. Both of our refactored BHR and NHR plasmid systems produced zones of clearing that were comparable in size to those produced by a control strain (SK01) with a previously engineered version of the two-plasmid MccV secretion system (9), demonstrating that the new refactored genetic parts and plasmid configuration support microcin secretion.

**Fig. 3:**
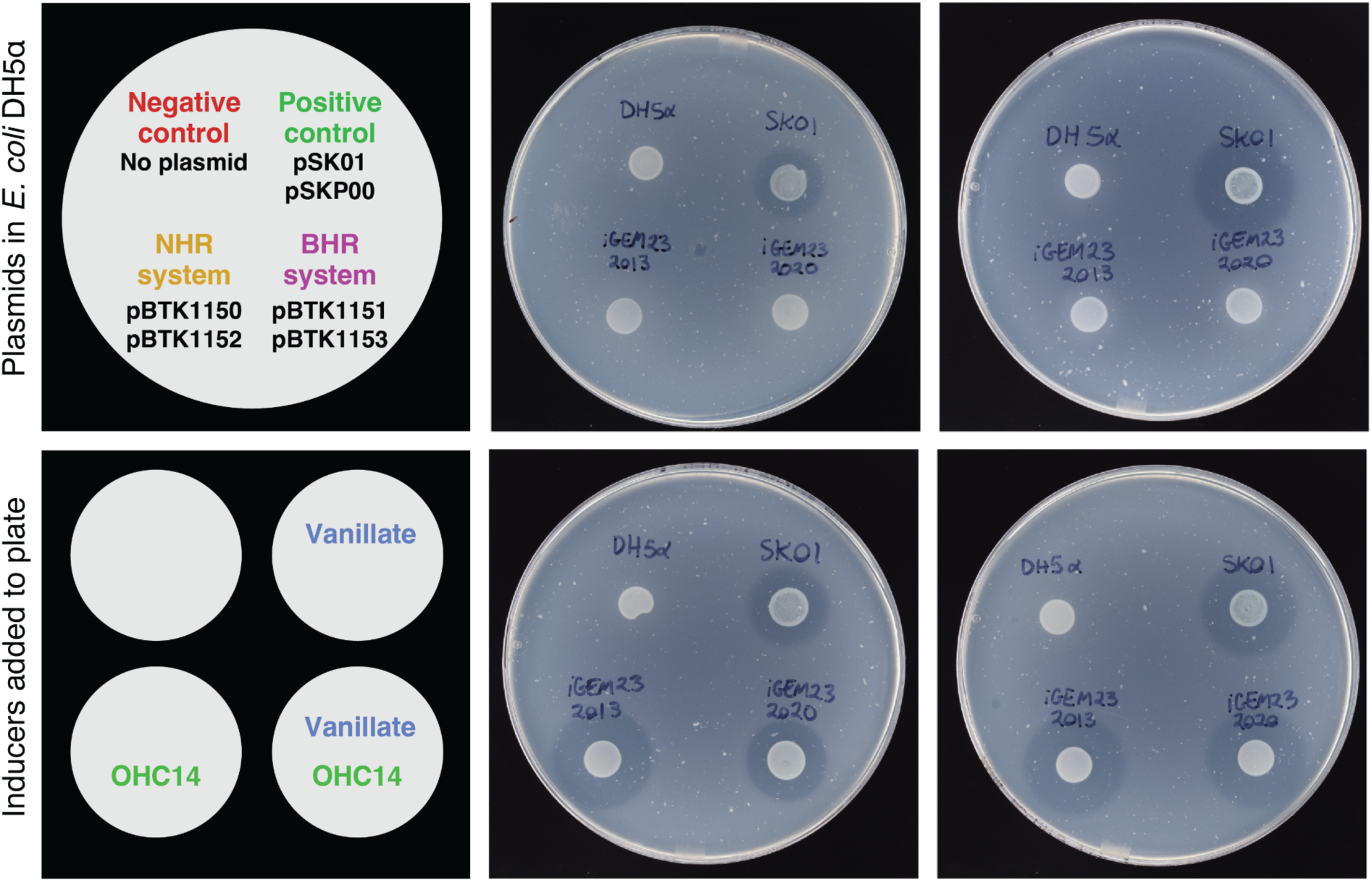
Two-plasmid secretion systems assembled from refactored T1SS genetic parts form zones of inhibition when MccV expression is induced. Each plate contains a top agar layer inoculated with microcin-susceptible strain *E. coli* W3110 that forms a turbid haze after incubation for growth. Four strains were spotted on each plate in the same arrangement: *E. coli* DH5α with no plasmids (negative control), *E. coli* W3110 with a previously validated two-plasmid secretion system (positive control) that is not under inducible control, or *E. coli* DH5α with either the narrow-host-range (NHR) or broad-host-range (BHR) two-plasmid systems we assembled from our refactored genetic parts. In the NHR and BHR systems, vanillate induces T1SS expression and OHC14 induces expression of MccV and its immunity protein. The upper left plate contains no inducers, the upper right plate contains vanillate only, the lower left plate contains OHC14 only, and the lower right plate contains both vanillate and OHC14.

### Plant-associated bacteria encode putative class II microcins

Next, we wanted to use our modular T1SS plasmids to explore the functions of novel microcins. Class II microcins have primarily been characterized in *E. coli* and its relatives within the *Enterobacteriaceae* family (2, 3). However, new bioinformatics approaches have recently led to the identification and characterization of microcins in other bacterial groups (2, 11, 31). We used the *cinful* bioinformatics pipeline (11) to identify putative class II microcins in the genomes of γ-proteobacteria in the genera *Pantoea* and *Erwinia* (*Erwiniaceae* family) and *Xanthomonas* (*Xanthomonadaceae* family). These groups mainly consist of plant-associated bacteria and include important plant pathogens. We manually filtered the output of *cinful* to a final list of 159 class II microcin candidates by applying several additional criteria (see **Methods**). The filtering procedure included eliminating candidates with a C-terminal sequence motif typically found in class IIb microcins in order to enrich for putative class IIa microcins in the final list. We constructed phylogenetic trees within each genus to cluster these sequences into similarity groups (**Fig. S1**). We selected 23 final microcin candidates representing different sequence clusters for experimental testing (**Fig. 4**). Putative microcins in *Pantoea* and *Erwinia* genomes often clustered together whereas microcins from *Xanthomonas* genomes were more distinct, possibly indicating that there is horizontal gene exchange of microcins within the *Erwiniaceae*.

**Fig. 4:**
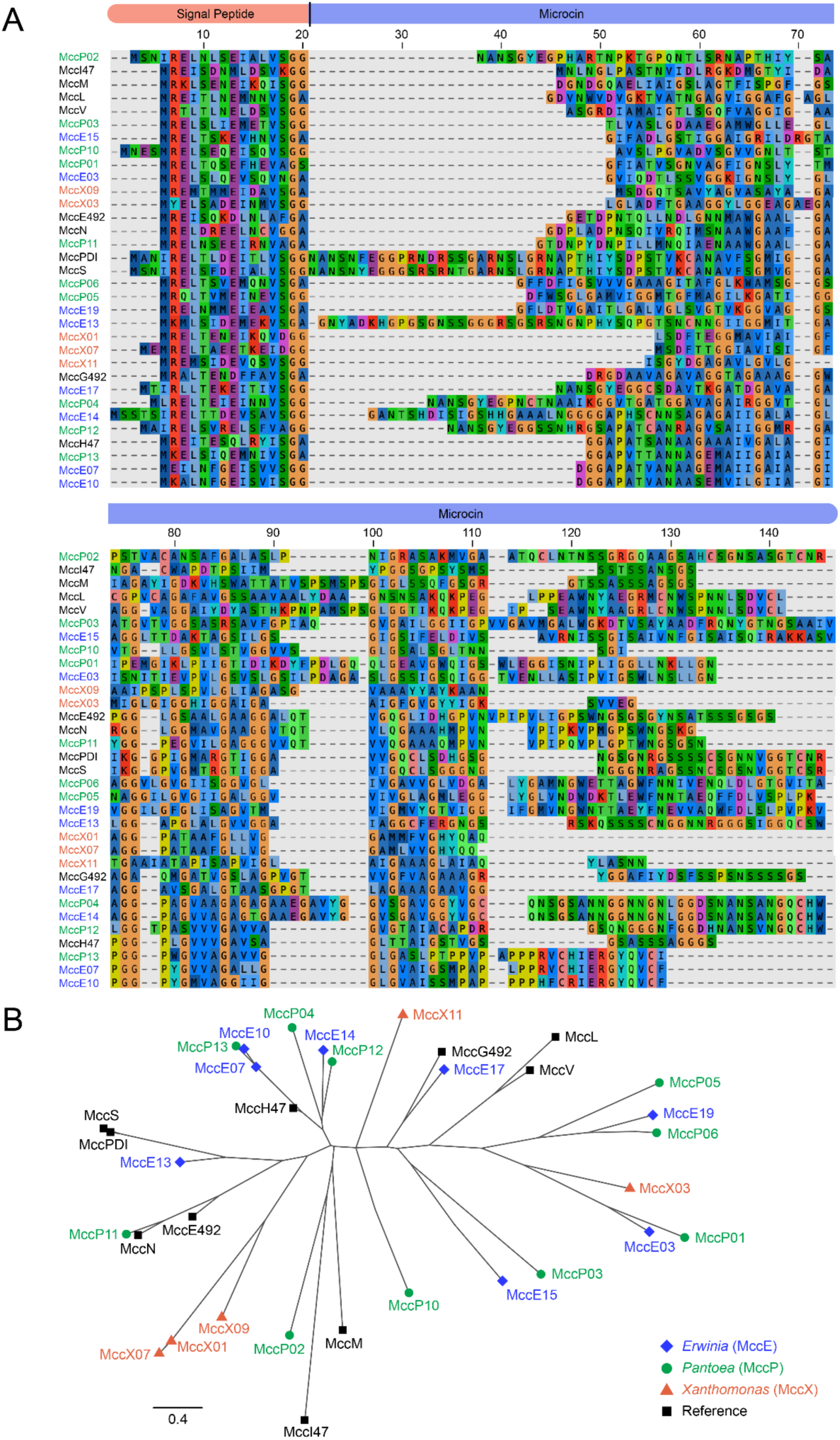
Microcin candidates identified in the genomes of bacteria from three plant-associated genera. **(A)** Multiple sequence alignment of 23 predicted class II microcins identified in *Pantoea* (MccP), *Erwinia* (MccE), and *Xanthomonas* (MccX) genomes that are representative of the full set of 191 unique candidates that were identified. Ten known class II microcins from *E. coli* and *Klebsiella* strains are included in the alignment for comparison. **(B)** Unrooted approximate maximum likelihood phylogenetic tree constructed from the mature microcin portion of the multiple sequence alignment (removing predicted signal peptides). The distance scale is based on amino acid sequence similarity using the BLOSUM45 matrix with a correction for multiple substitutions.

### Expression of class II microcin candidates inhibits *E. coli* growth

The antibacterial activity of some class II microcins can be detected using a self-inhibition assay (2). If a microcin-secreting strain lacks the cognate immunity protein and is susceptible to that microcin, induction of microcin expression will slow its growth and/or lead to self-killing that delays or prevents a culture from becoming fully turbid. We cloned each of the 23 microcin candidates identified in the genomes of bacteria from plant-associated genera into the two-plasmid NHR secretion system. Then, we monitored the turbidity of cultures of *E. coli* W3110 strains transformed with these plasmids when microcin expression was induced. Uninduced cultures of each strain were grown in parallel to serve as a reference point for when the putative microcin is not expressed and there should not be any inhibition. A strain with MccV cloned into the two-plasmid NHR secretion system without its immunity protein was included as a positive control. A strain with a random peptide sequence (G3P2), which does not exhibit inhibition in this assay (9), cloned into the NHR system was included as a negative control.

Two trials of the self-inhibition assay were conducted for each of these 25 strains (**Fig. 5, Fig. S2**). We tested whether induction of microcin expression caused a significant decrease in turbidity at the 4- and 12-hour timepoints (one-tailed *t*-tests, Benjamini-Hochberg adj. *p* < 0.05 and ≥10% mean reduction in OD600). Seven candidates (MccP04, MccP11, MccP12, MccP13, MccE07, MccE14, and MccX09) exhibited significantly reduced turbidity associated with microcin induction at either the 4- or 12-hour timepoint in both trials. Induction of MccP11, MccP12, MccP13, MccE07, or MccE14 expression caused the increase in turbidity to slow around 2-3 hours and appeared to prevent cultures from reaching full saturation. Induction of MccP04 and MccX09 expression only had an effect after 4 hours, although the MccP04 caused a large reduction in turbidity at 12 hours. Four other microcin candidates (MccP03, MccP06, MccP10, and MccE17) exhibited effects on growth that were only significant in one of the two trials. Most also diverged from the uninduced treatment 6 or more hours into the experiment.

**Fig. 5:**
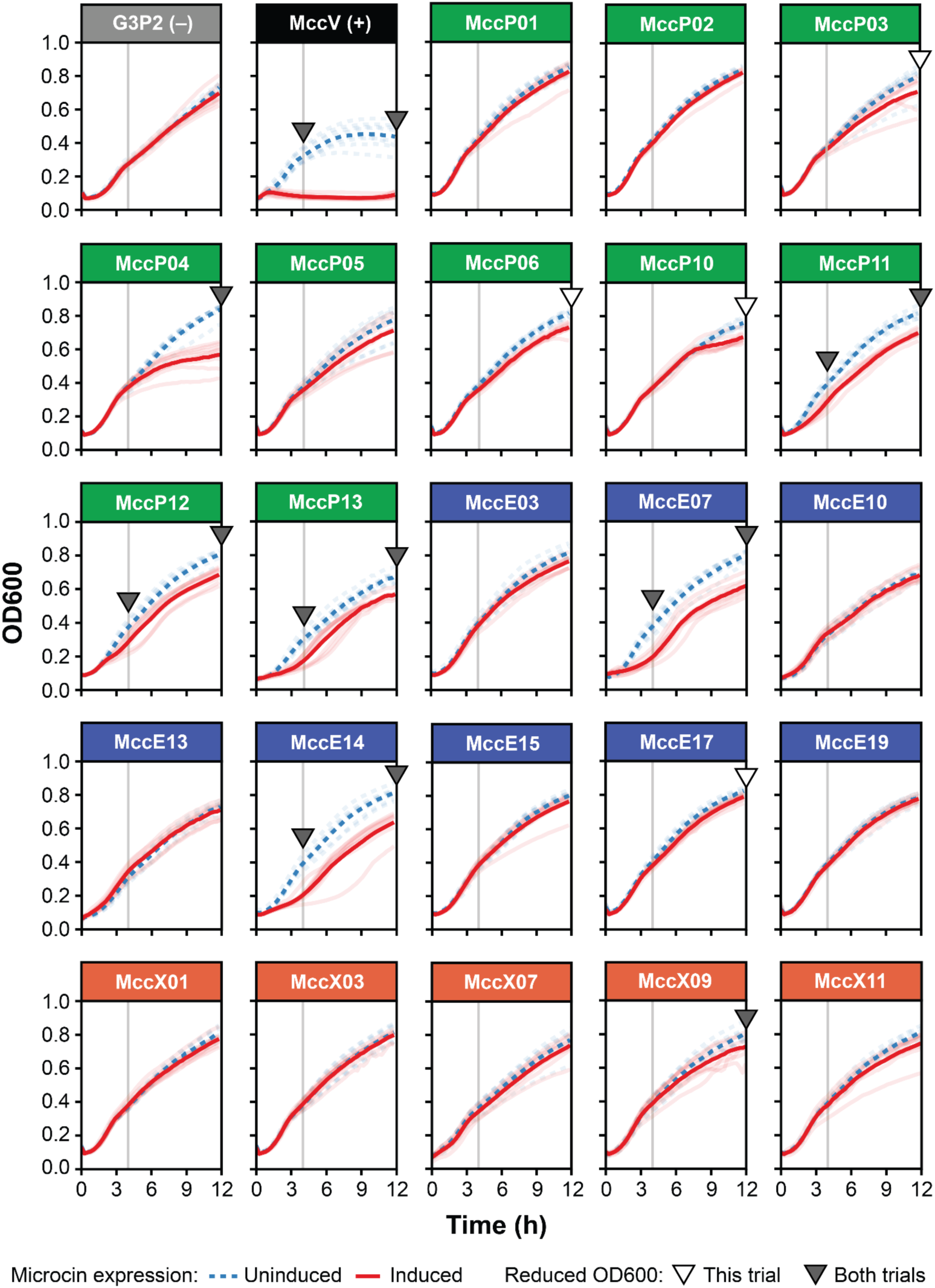
Some microcin candidates exhibit antibacterial activity in *E. coli* self-inhibition assays. Growth curves of *E. coli* W3110 strains containing different microcin candidates cloned into the two-plasmid secretion system were collected with (red solid lines) or without (blue dashed lines) OHC14 induction. The most saturated lines in each panel show the mean of eight biological replicates shown as transparent lines. Triangles indicate a significant difference between the OD600 values of the induced and uninduced cultures at 4 h or 12 h in both trials of this experiment (filled) or only in the trial shown in this figure (unfilled) (see **Methods**). Analogous plots for the second trial are shown in **Figure S2**.

Overall, these results suggest that some of the microcin candidates identified in plant-associated bacteria exhibit some degree of antibacterial activity against *E. coli* W3110, though none has as strong of an effect as the *E. coli*-derived microcin MccV used as a positive control..

## Discussion

We refactored the *E. coli* MccV T1SS into interchangeable part modules compatible with the Golden Gate assembly scheme used by the BTK and YTK synthetic biology toolkits (14, 15). The vanR and cinR inducible promoter systems that we refactored and validated also add missing functionality to these toolkits. Other BTK promoter parts (22, 26), plasmid backbones (27), and integration cassettes (21), which have been used to engineer a variety of bacterial species in prior work, can be combined with our new parts to create tailored secretion systems. This modular approach makes it straightforward to optimize and adapt microcin T1SSs for new applications. For example, one could combine all T1SS and cargo elements into a single plasmid or integration cassette via Stage 2 assembly (15) or refactor the operons and gene clusters needed for post-translationally modifying different class IIb microcins into new parts (16). Because different cargos, including diverse proteins <100 amino acids, have been shown to be effectively secreted by the *E. coli* MccV T1SS (9), there are many possible applications beyond microcins.

We identified many putative class II microcins in the genomes of plant-associated *Pantoea*, *Erwinia*, and *Xanthomonas* strains and used our modular two-plasmid system to test them for activity against *E. coli*. With the exception of the predicted class IIb microcin *Ps* G492 (16), these are the first class II microcins from these bacterial groups that have been cloned and experimentally screened for antibacterial activity to our knowledge. These groups include bacteria that are plant pathogens, such as *Erwinia amylovora* (32) and *Pantoea ananatis* (33) strains. They also include plant commensals and even biocontrol strains that are applied to protect crops from related pathogens. For example, *Pantoea vagans* C9-1 is used against fire blight caused by *E. amylovora* (34). We identified microcin candidates with closely related and even identical sequences in *Pantoea* and *Erwinia* strains, which could indicate that microcin T1SS gene clusters are often horizontally transferred between *Erwiniaceae* family bacteria. By contrast, the sequences of microcin candidates found in *Xanthomonas* genomes tended to cluster separately. Class IIb microcins were recently identified in the genomes of plant pathogens from genera other than those examined here (16). Overall, these results suggest that microcins may be important for mediating competition within plant microbiomes.

None of the 23 putative microcins we tested exhibited antibacterial activity against *E. coli* that was comparable in strength to MccV in our self-inhibition assays. The limited inhibition that we did observe for some candidates only delayed growth or affected cultures as they neared stationary phase. These results could reflect aspects of microcin biology or technical limitations. Because many class II microcins are thought to have evolved to target close relatives (1, 35), it is likely that *E. coli* is not susceptible or as susceptible to these microcins as their species of origin would be. *E. coli* may lack a necessary receptor for entry into cells or an optimal target for inhibition, for example. Furthermore, while MccV exhibits bactericidal activity (36, 37), other class II microcins only exhibit bacteriostatic activity (17). Bacteriostatic activity is compatible with how expression of some microcin candidates affected *E. coli*, particularly if their receptors or targets were not expressed until later in growth. Finally, even though we sought to limit the candidates we tested to class IIa microcins, some may require post-translational modifications for full activity, and we did not attempt to include any operons encoding these activities in our constructs. On the technical side, it is possible that the *E. coli* MccV T1SS does not support efficient expression and/or secretion of the heterologous microcins. Our results could also potentially be explained if inducing expression of these microcin candidates in *E. coli* led to a nonspecific burden (38, 39) rather than growth arrest due to a specific inhibitory interaction.

In the future, our modular system could be used to further characterize these and other microcins in ways that address these potential shortcomings. Deploying these systems using plasmids or integration vectors that function in *Pantoea*, *Erwinia*, and *Xanthomonas* strains, such as the two-plasmid BHR system we constructed, would make it possible to perform self-inhibition assays in more closely related hosts. Cloning MFP and PCAT proteins, signal peptides, immunity proteins, and candidate post-translational tailoring factors as genetic parts could be used to better implement heterologous secretion from *E. coli* or other species. In particular, adding candidate immunity proteins in our constructs would make it possible to test whether these rescue the self-inhibition that we observed and then to see if *E. coli* secreting the microcin candidate could inhibit plant pathogens either in the context of zone of inhibition assays or co-culture competition assays that can be more sensitive for detecting bacteriostatic effects.

More generally, secretion of microcins can be used to inhibit pathogens and potentially for microbiome engineering applications. For example, microcins could be mined from related *Pantoea* and *Erwinia* strains and added to the factors produced by *P. vagans* C9-1 in order to make it a more effective biocontrol agent. Editing the microcins encoded in a strain might also be key to allowing it to invade and establish in a microbiota, as has been explored with colicins and other secreted effectors (40, 41). Many class II microcins predicted in genomes remain to be characterized, and some microcins may have effects on plant or animal hosts in addition to or rather than targeting bacteria (1, 6). Microcin secretion systems have also been used to produce heterologous bioactive peptides that affect animal hosts (9), proving that this mode of action is possible. Thus, our work contributes to continuing efforts to characterize and adapt both microcins and their versatile secretion systems for a variety of scientific, engineering, medical, and agricultural goals.

## Data availability

All data generated or analyzed during this study are included in this published article and its supplementary information files.

## Abbreviations

BHR: Broad-host-range
BTK: Bee Microbiome Toolkit
Cam: Chloramphenicol
Carb: Carbenicillin
GGA: Golden Gate assembly
Kan: Kanamycin
MFP: Membrane fusion protein
NHR: Narrow-host-range
OD600: Optical density at 600 nm
OHC14: 3-hydroxytetradecanoyl-homoserine lactone
PCAT: Peptidase-containing ABC transporter
RFU: Relative fluorescence units
T1SS: Type I secretion system
YTK: Yeast Toolkit

## Acknowledgments

We thank other members of The University of Texas at Austin 2023 iGEM Team for their help with project ideation and initial experiments: Aria Welch, Aurea Le, Elizabeth Manriquez, Ivy Huynh, Meghna Vergis, Nate Brant, Neha Donthineni, Samer Salman, Sneha Chandak, Sreya Das, and Stephanie Sustaita. We thank Sun-Young Kim and Bryan Davies for providing strains and plasmids and Peng Geng, Shaunak Kar, and Victor Li for creating entry and dropout vectors. We acknowledge the Texas Advanced Computing Center (TACC) at The University of Texas at Austin for providing high-performance computing resources.

## Contributions

Conceptualization: AKM, KC, BDM, VBI, KED, JKP, DMM, and JEB; Investigation: AKM, KC, BDM, VBI, and KED; Reagents: JKP and AJV; Supervision: AJV, JKP, DMM, JEB; Visualization: AKM, KC, BDM, VBI, JEB; Formal analysis: AKM, KC, BDM, VBI, and JEB; Writing – original draft: AKM, KC, BDM, VBI, KED, JEB; Writing – review & editing: AKM, KC, BDM, VBI, KED, AJV, JKP, DMM, JEB.

## Ethics declarations

### Ethics approval and consent to participate

Not applicable.

### Consent for publication

Not applicable.

## Supplementary Information

**Table S1.** Strains used in this work.

**Table S2.** Plasmids used in this work.

**Table S3.** Overhangs defining part types in the Golden Gate assembly scheme.

**Table S4.** Final microcin candidates.

**Fig S1.** Passed microcin candidate phylogenetic trees by genus.

**Fig S2.** Second trial of the *E. coli* self-inhibition assays.

**Data S1.** Plasmid genbank files.

**Data S2.** Unprocessed *cinful* output files and microcin candidate lists, alignments, and phylogenetic trees.

