## Supplementary Figures for "Refactored genetic parts for modular assembly of the *E. coli* MccV type I secretion system used to screen class II microcin candidates from plant-associated bacteria"

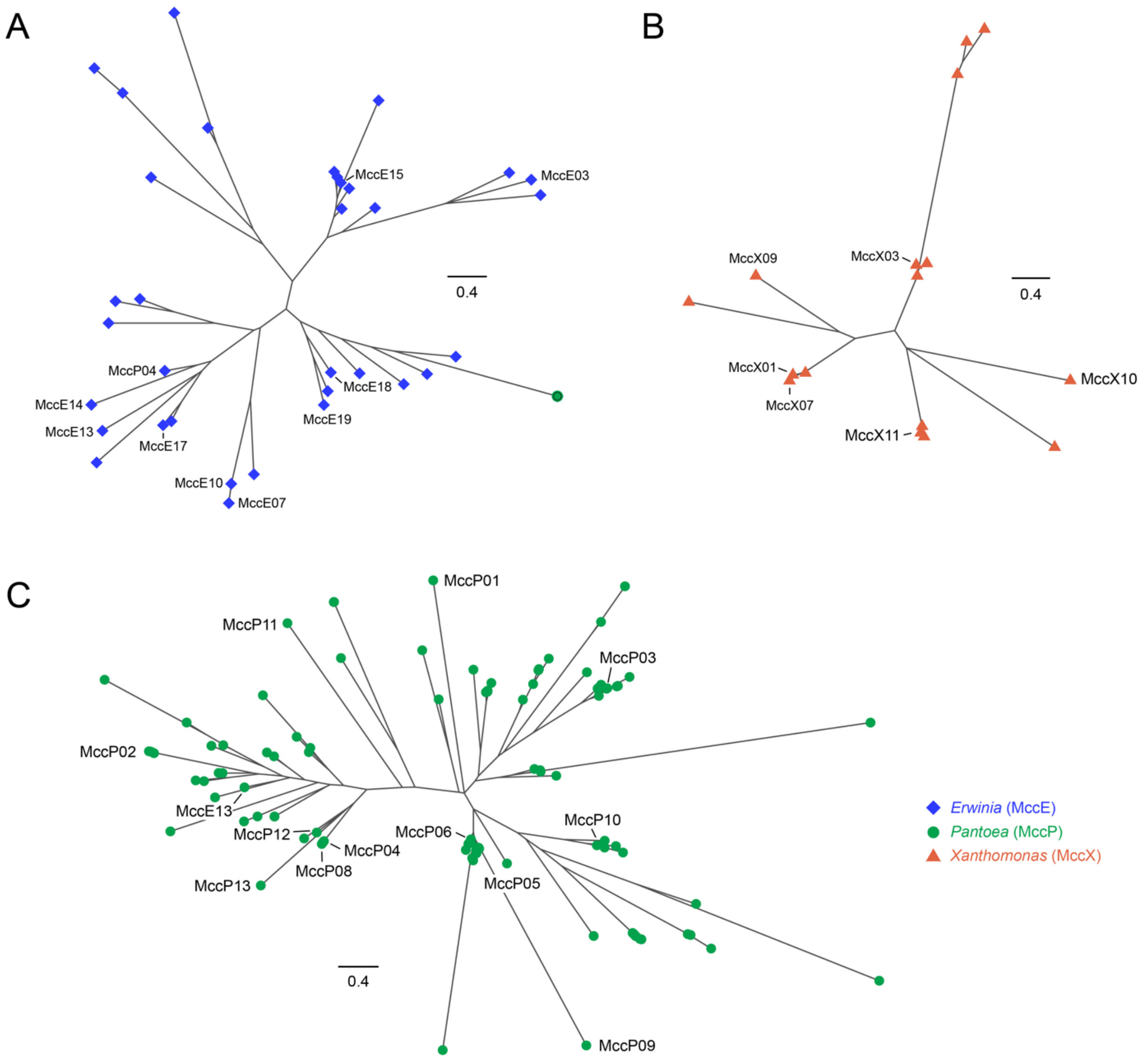


**Fig. S1:** Passed microcin candidate phylogenetic trees by genus. (A) *Erwinia* microcin candidates. (B) *Xanthomonas* microcin candidates. (C) *Pantoea* microcin candidates. For all panels, unrooted approximate maximum likelihood phylogenetic trees were constructed from the mature microcin portion of the multiple sequence alignment (removing predicted signal peptides). The distance scales are based on amino acid sequence similarity using the BLOSUM45 matrix with a correction for multiple substitutions.

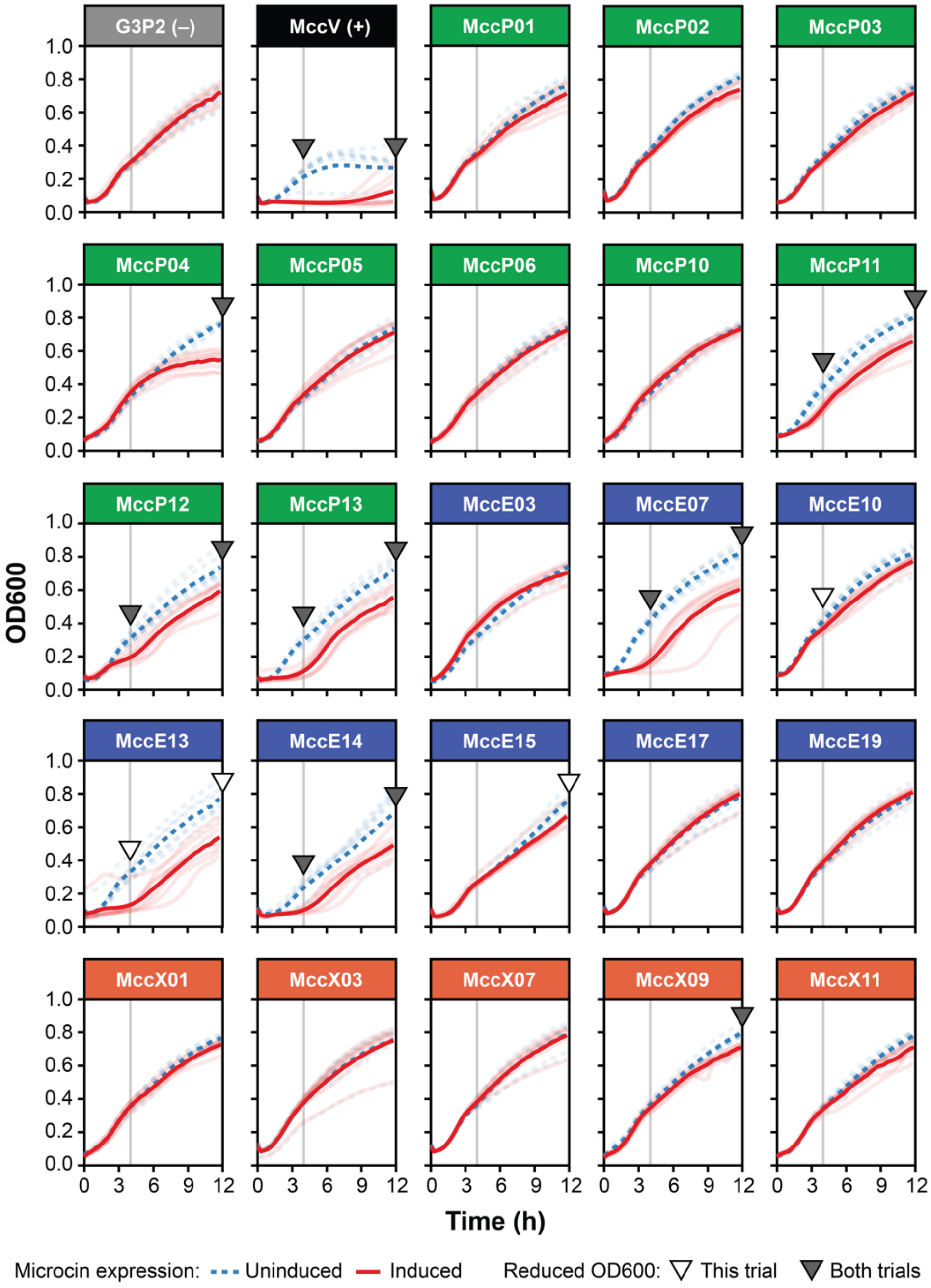


**Fig. S2:** Some microcin candidates exhibit antibacterial activity in *E. coli* self-inhibition assays. Growth curves of *E. coli* W3110 strains containing different microcin candidates cloned into the two-plasmid secretion system were collected with (red solid lines) or without (blue dashed lines) OHC14 induction. The most saturated lines in each panel show the mean of eight biological replicates shown as transparent lines. Triangles indicate a significant difference between the OD600 values of the induced and uninduced cultures at 4 h or 12 h in both trials of this experiment (filled) or only in the trial shown in this figure (unfilled) (see **Methods**). Analogous plots for the first trial are shown in **Figure 5**.
